# Residue-by-residue analysis of cotranslational membrane protein integration *in vivo*

**DOI:** 10.1101/2020.09.27.315283

**Authors:** Felix Nicolaus, Ane Metola, Daphne Mermans, Amanda Liljenström, Ajda Krč, Salmo Mohammed Abdullahi, Matthew Zimmer, Thomas F. Miller, Gunnar von Heijne

**Affiliations:** Department of Biochemistry and Biophysics, Stockholm University, SE-106 91 Stockholm, Sweden; Faculty of Chemistry and Chemical Technology, University of Ljubljana, Večna pot 113, 1000 Ljubljana, Slovenia; Science for Life Laboratory Stockholm University, Box 1031, SE-171 21 Solna, Sweden; California Institute of Technology, Division of Chemistry and Chemical Engineering, Pasadena, CA 91125, USA

**Keywords:** cotranslational protein folding, membrane protein, BtuC, GlpG, EmrE

## Abstract

We follow the cotranslational biosynthesis of three multi-spanning *E. coli* inner membrane proteins *in vivo* using high-resolution Force Profile Analysis. The force profiles show that the nascent chain is subjected to rapidly varying pulling forces during translation, and reveal unexpected complexities in the membrane integration process. We find that an N-terminal cytoplasmic domains can fold in the ribosome exit tunnel before membrane integration starts, that charged residues and membrane-interacting segments such as re-entrant loops and surface helices flanking a transmembrane helix (TMH) can advance or delay membrane integration, and that point mutations in an upstream TMH can affect the pulling forces generated by downstream TMHs in a highly position-dependent manner, suggestive of residue-specific interactions between TMHs during the integration process.

---

Most integral membrane proteins are cotranslationally integrated into their target membrane with the help of translocons such as the bacterial SecYEG/YidC complex and the eukaryotic Sec61 complex (*1*). While the energetics of translocon-mediated integration of transmembrane α-helices (TMHs) is reasonably well understood (*2*), the actual integration process is not, other than in general terms. We have shown that Force Profile Analysis (FPA) – a method in which a translational arrest peptide (AP) engineered into a target protein serves as a sensor to measure the force exerted on a nascent polypeptide chain during translation – can be used to follow the cotranslational folding of soluble proteins and the membrane integration of a model TMH (*3-5*). Here, we have applied FPA to follow the cotranslational membrane integration of three multi-spanning *E*.*coli* inner membrane proteins (EmrE, GlpG, BtuC), providing the first residue-by-residue picture of membrane protein integration *in vivo*.

FPA takes advantage of the ability of APs to bind in the upper parts of the ribosome exit tunnel and thereby pause translation on a specific codon (*6*). The duration of the pause is reduced in proportion to pulling forces exerted on the nascent chain from outside the ribosome (*7, 8*), i.e., APs can act as force sensors, and can be tuned by mutation to react to different force levels (*9*). In an FPA experiment, a series of constructs is made in which a force-generating sequence element (e.g., a TMH) is placed an increasing number of residues *N* away from an AP, which in turn is followed by a C-terminal tail, Fig. 1a. In constructs where the TMH engages in an interaction that generates a pulling force on the nascent chain at the point when the ribosome reaches the last codon of the AP, pausing will be prevented and mostly full-length protein will be produced during a short pulse with [^35^S]-Met, Fig. 1b (middle). In contrast, in constructs where little force is exerted on the AP, pausing will be efficient and more of the arrested form of the protein will be produced. The fraction full-length protein produced, *f*_*FL*_=*I*_*FL*_/(*I*_*FL*_+*I*_*A*_), where *I*_*FL*_ and *I*_*A*_ are the intensities of the bands representing the full-length and arrested species on an SDS-PAGE gel, Fig. 1c, can therefore be used as a proxy for the force exerted on the AP in a given construct (*8, 10, 11*). A plot of *f*_*FL*_ vs. *N* – a force profile (FP) – thus can provide a detailed picture of the cotranslational process in question, as reflected in the variation in the force exerted on the AP during translation.

**Figure 1.**
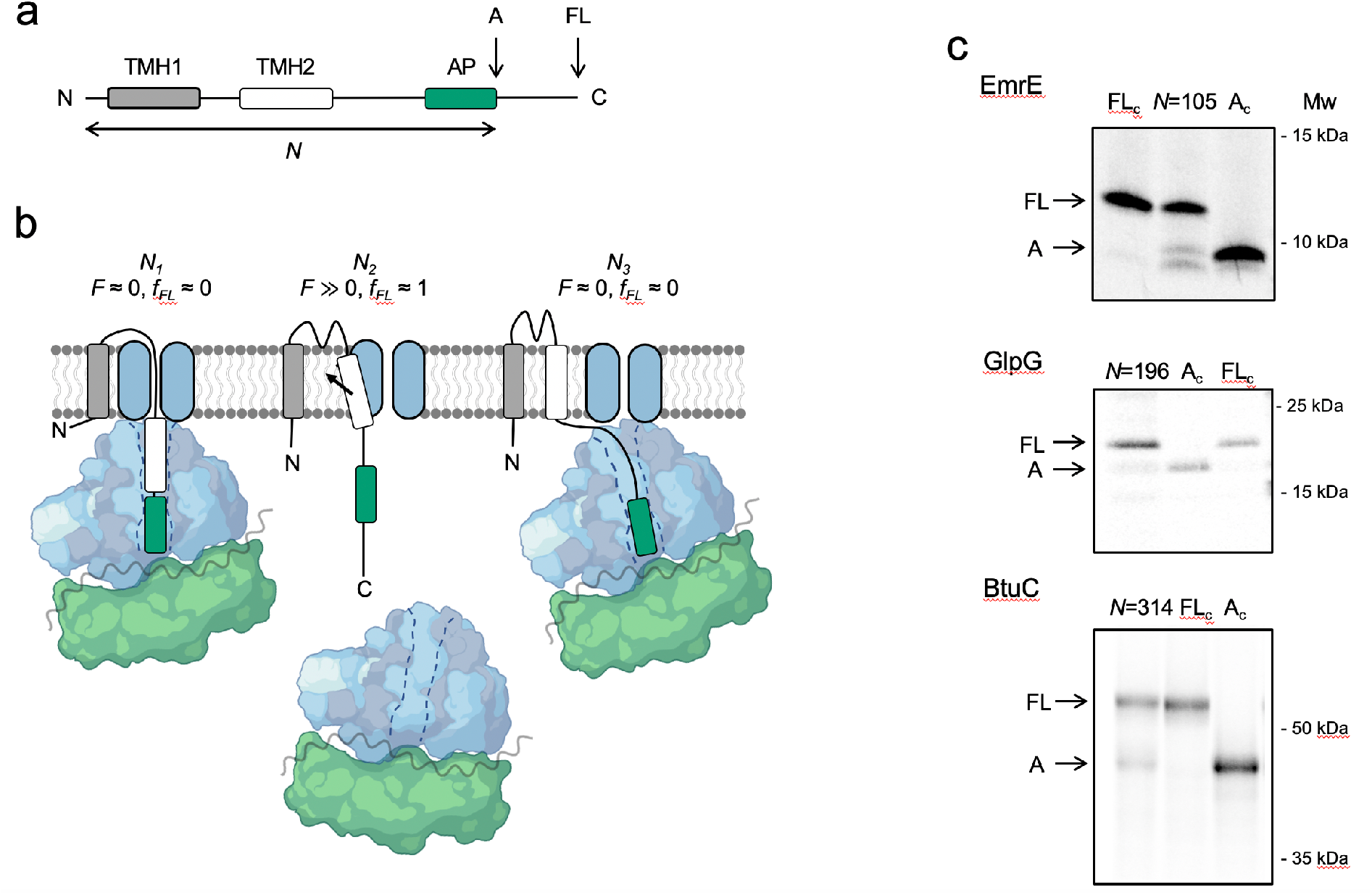
The force profile assay. (a) Basic construct. Arrested (*A*) and full-length (*FL*) products are indicated. (b) At construct length *N*_*1*_, TMH2 has not yet entered the SecYEG channel and no pulling force *F* is generated. At *N*_*2*_, TMH2 is integrating into the membrane and *F* ≫0. At *N*_*3*_, TMH2 is already integrated and *F*≈0. (c) SDS-PAGE gels showing *A* and *FL* products for EmrE(C_out_)(*N*=105), GlpG(*N*=196), and BtuC(*N*=314). Control constructs *A*_*C*_ and *FL*_*c*_ have, respectively, a stop and an inactivating Ala codon replacing the last Pro codon in the AP.

We chose EmrE as an example of a small, relatively simple 4-TMH protein. EmrE is a dual-topology protein, i.e., the monomers integrate into the inner membrane in a 50-50 mixture of N_in_-C_in_ and N_out_-C_out_ topologies; two oppositely oriented monomers then assemble into an antiparallel dimer (*12, 13*). To avoid potential complications caused by the dual topology, we used EmrE(C_out_), a mutant version that adopts the N_out_-C_out_ topology (*13*), and the relatively weak SecM(*Ec*) AP (*4*), Fig. 2a. A series of EmrE(C_out_)-AP constructs (see Table S1 for sequences) was used to obtain the FP shown in Fig. 2b (orange curve), at 2-5 residues resolution. Also shown is a FP derived from a coarse-grained molecular dynamics simulation (CGMD-FP, grey) (*14*); a hydrophobicity plot (HP) is included in Fig. S1.

**Figure 2.**
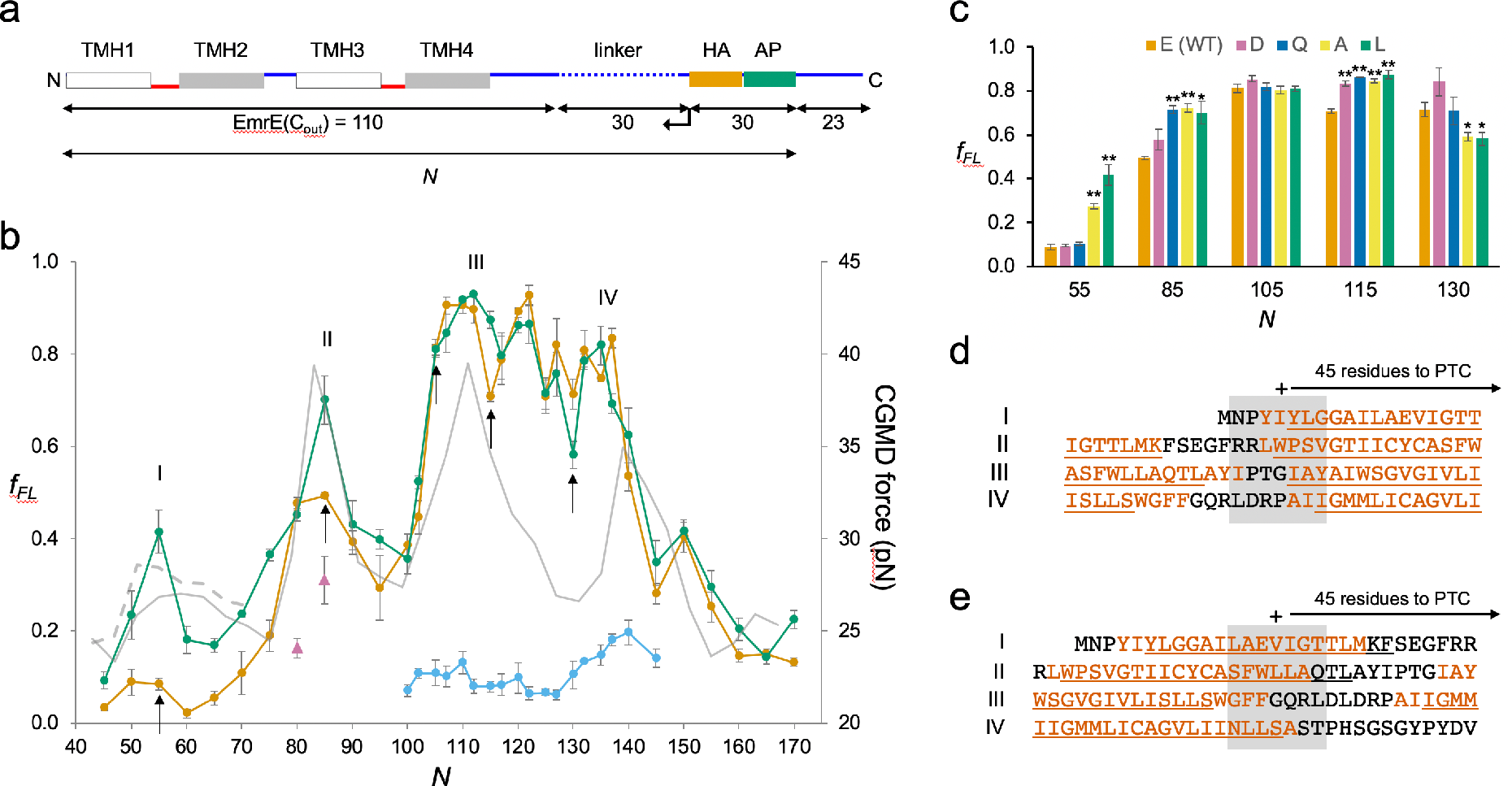
EmrE(C_out_). (a) Construct design. EmrE(C_out_) is shortened from the C-terminal end of the LepB-derived linker (dotted), as indicated by the arrow. Cytoplasmic (red) and periplasmic (blue) loops, and lengths of full-length EmrE(C_out_), LepB-derived linker, HA tag+AP, and C-terminal tail are indicated. Since the 30-residue HA+AP segment is constant in all constructs, the FP reflects nascent chain interactions occurring outside the ribosome exit tunnel. (b) FPs for EmrE(C_out_) (orange), EmrE(C_out_,E^14^L) (green), EmrE(C_out_) with SecM(*Ec*-sup1) AP (blue), EmrE(C_out_, I^37^I^38^→NN) (magenta triangles), and CGMD-FP calculated with a −100 mV membrane potential (grey). (c) Effects of mutations in E^14^ on *f*_*FL*_ values for the *N* values indicated by arrows in panel *b. p* values (2-sided t-test): *p* < 0.05, *; *p* < 0.01, **. (d-e) Sequences corresponding to peaks I-IV aligned from their *N*_*start*_ (d) and *N*_*end*_ (e) values. + sign indicates 45 residues from the PTC. Hydrophobic TMH segments shown in orange and membrane-embedded α-helices underlined (PDB ID: 3B5D). Error bars in panels *b* and *c* indicate SEM values.

We have previously shown that a model TMH composed of Ala and Leu residues generates a peak in the FP that reaches half-maximal amplitude (*N*_*start*_) when the N-terminal end of the TMH is ∼45 residues away from the polypeptide transferase center (PTC) (*4*), and a real-time FRET study of cotranslational membrane integration found that the N-terminal ends of N-terminal TMHs reach the vicinity of the SecYEG translocon when they are 40-50 resides away from the PTC (*15*). For EmrE(C_out_) TMH1, this would correspond to constructs with *N*≈50. However, the *f*_*FL*_ values are hardly above background in this region of the FP. Due to the functionally important E^14^ residue, TMH1 is only marginally hydrophobic and does not become firmly embedded in the membrane until the protein dimerizes (*16*). To ascertain whether the lack of a peak in the FP corresponding to the membrane integration of TMH1 is because of its low hydrophobicity, we mutated E^14^ to L. Indeed, in the FP obtained for EmrE(C_out_,E^14^L), Fig. 2b (green curve), a clear peak appears at the expected chain length *N*_*start*_≈50 residues. Mutation E^14^A yields an *f*_*FL*_ value intermediate between EmrE(C_out_,E^14^L) and EmrE(C_out_) at *N*=55, Fig. 2c, while *f*_*FL*_ for EmrE(C_out_,E^14^D) and EmrE(C_out_,E^14^Q) are the same as for EmrE(C_out_).

Peak II corresponds to a situation where the N-terminal end of TMH2 is ∼45 residues from the PTC, Fig. 2d. The double mutation I^37^I^38^→NN in TMH2 reduces *f*_*FL*_ at *N*=80 and 85 (magenta triangles), as expected. Unexpectedly, however, the E^14^L, E^14^A, and E^14^Q (but not the E^14^D) mutations in TMH1 increase *f*_*FL*_ at *N*=85, Fig. 2c, showing that a negatively charged residue in position 14 in TMH1 specifically reduces the pulling force generated by TMH2 at *N*=85, i.e., when about one-half of TMH2 has integrated into the membrane. Likewise, *f*_*FL*_ values at *N*=115 and 130 (but not at *N*=105, included as a negative control) are specifically affected by mutations in E^14^: at *N*=115 (one-half of TMH3 integrated), all four mutations in position 14 increase *f*_*FL*_ relative to E^14^, while at *N*=130 (beginning of TMH4 integration), the E^14^A and E^14^L mutations decrease *f*_*FL*_, Fig. 2c. FPA thus reveals long-range effects of mutations in TMH1 on three specific steps in the membrane integration of downstream TMHs.

Peak III has *N*_*star*t_≈102 residues, with the N-terminal end of TMH3 ∼45 residues from the PTC, Fig. 2d. Peak IV is difficult to locate precisely in the FP, but is seen at *N*_*star*t_≈132 residues when the strong SecM(*Ec-sup1*) AP (*17*) is used (blue curve), again with the N-terminal end of TMH4 ∼45 residues from the PTC, Fig. 2d. As shown in Fig. 2e, the TMHs cease generating a pulling force when their C-terminal ends are ∼45 residues away from the PTC, indicating that they are fully integrated at this point.

We next studied GlpG, a monomeric 6-TMH rhomboid protease with a cytoplasmic N-terminal domain (NTD) (*18, 19*), Fig. 3a, a protein that allows us to follow the cotranslational folding of a soluble domain and integration of a membrane domain in the same experiment. The FP is shown in Fig. 3b (orange curve). For unknown reasons, constructs with *N*≈140-200 residues gave rise to multiple bands on the gel that were difficult to interpret; this problem did not arise when a portion of the LepB protein was appended at the N-terminus, Fig. 3a, and the corresponding *f*_*FL*_ values are shown in the FP (*N*=131-224). The FP was obtained at 5-residue resolution, except for the portion *N*=178-215 which we measured with single-residue resolution. *N*_*start*_ and *N*_*end*_ values for peaks II-VII are indicated in Fig. S2b,c.

**Figure 3.**
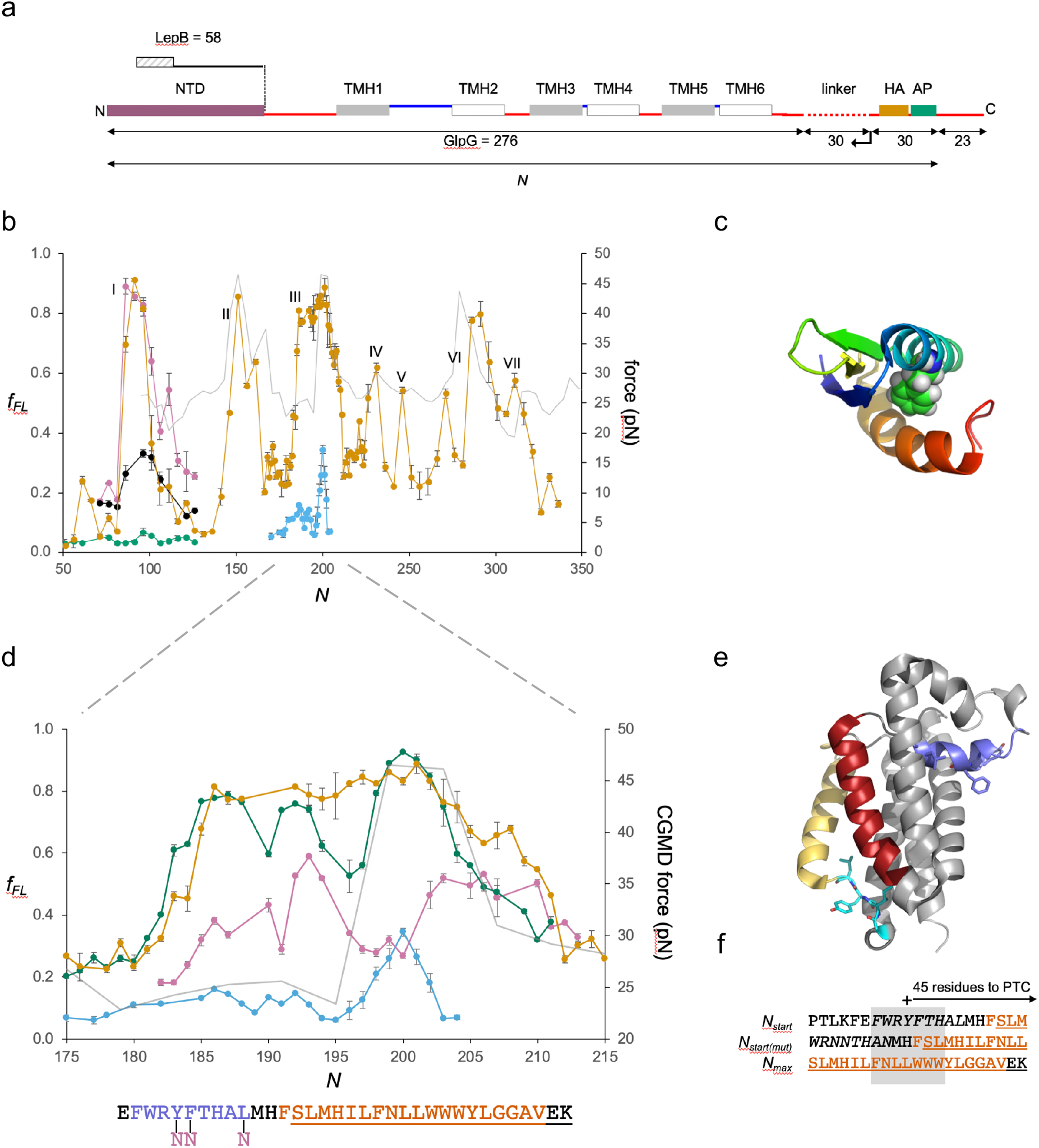
GplG. (a) Construct design, c.f., Fig. 2a. The N-terminal LepB fusion is indicated. (b) FPs for GlpG and LepB-GlpG (*N*=131-224) (orange), NTD(F^16^E) (green), *in-vitro* translated NTD (magenta) and NTD(F^16^E) (black), GlpG with SecM(*Ec*-Sup1) AP (blue), and CGMD-FP calculated with a −100 mV membrane potential (grey). Error bars indicate SEM values. (c) NTD (PDB ID: 2LEP), with F^16^ in spacefill. (d) Enlarged FPs for LepB-GlpG with SecM(*Ec*) AP (orange), SecM(*Ec-Ms*) AP (green), SecM(*Ec*-sup1) AP (blue), and GlpG(Y^138^F^139^L^143^→NNN) with SecM(*Ec-Ms*) AP (magenta). CGMD-FP in grey. (e) Structure of GlpG with the periplasmic surface helix in blue, TMH2 in red, the membrane-associated cytoplasmic segment in cyan, and TMH5 in yellow. Y^138^F^139^L^143^ and G^222^I^223^Y^224^L^225^ are shown as sticks. (f) Peak III sequences aligned from their *N*_*start*_ and *N*_*max*_ values for GlpG and mutant GlpG(Y^138^F^139^L^143^→NNN). The surface helix is italicized.

Peak I, at *N*_*start*_≈84 residues, is conspicuously close to what would be expected from previous studies of cotranslational folding of small globular domains in the ribosome exit tunnel (*5*). To verify that the peak represents folding of the NTD, we recorded a FP for the NTD by *in vitro* transcription-translation in the PURE system (*20*), and further made a destabilizing point mutation (F^16^E) in the core of the NTD, Fig. 2c. The FP obtained *in vitro* (magenta) overlaps peak I in the *in vivo* FP, and the mutation strongly reduces the *in vivo* (green) and *in vitro* (black) *f*_*FL*_ values for peak I. Thus, the NTD folds inside the ribosome exit tunnel when its C-terminal end is ∼25 residues from the PTC, well before synthesis of the membrane domain has commenced.

Peaks II-VII in the FP correspond reasonably well to the CGMD-FP (grey) and HP (Fig. S2a). The unexpectedly low *N*_*start*_ value for peak III is caused by an upstream periplasmic surface helix (see below). Likewise, peak VI-a likely reflects the membrane integration of a hydrophobic, membrane-associated cytoplasmic segment located just upstream of TMH5, Fig. 3f and Fig. S2b, while peak VI-b represents the integration of TMH5 itself. In contrast, the unexpectedly high *N*_*start*_ value for peak IV indicates that integration of TMH3 commences only when its N-terminal end is ∼52 residues away from the PTC, possibly because of the tight spacing between TMH2 and TMH3.

As peak III saturates at *f*_*FL*_≈0.9 over a rather wide range, we sought a more detailed view by using the strong SecM(*Ec-*Sup1) AP (*17*), Fig. 3b,d (blue), and the medium-strong SecM(*Ec-Ms*) AP (*21*), Fig. 3d (green). The SecM(*Ec-*Sup1) FP allows a precise determination of *N*_*max*_=200 residues, at which point the middle of TMH2 (L^155^) is located 45 residues from the PTC, Fig. 3f. The SecM(*Ec-Ms*) FP reveals additional detail, with local minima at *N*≈190 and 195 corresponding roughly to when the polar residues H^145^ and H^150^ reach 45 residues from the PTC. We further recorded a SecM(*Ec-Ms*) FP (magenta) for the triple mutation Y^138^F^139^L^143^→NNN (Fig. 3e) that renders the periplasmic surface helix hydrophilic: the mutation shifts *N*_*start*_ for peak III from 182 to 190, with the N-terminal end of TMH2 ∼45 residues from the PTC, Fig. 3d,f. The triple mutation also widens the local minimum at *N=*195 up to *N*=201, shifts *N*_*max*_ from 200 to ∼204, and overall reduces the amplitude of peak III. Thus, the periplasmic surface helix engages in hydrophobic interactions already during its passage through the translocon – presumably by sliding along a partly open lateral gate (*22*) – and has multiple effects on the membrane integration of TMH2.

Finally, we studied BtuC, a vitamin B12 transporter with 10 TMHs, as an example of a large, multi-spanning protein with a complex fold (*23*). In order to improve expression, we added the N-terminal part of LepB to the BtuC constructs, Fig. 4a; constructs that we could measure without the LepB fusion gave similar *f*_*FL*_ values as those seen for the LepB fusions (Fig. S3b). We identified 11 peaks in the FP, Fig. 4b (orange), one more than could be accounted for by the 10 TMHs. Since it was not possible to provide an unequivocal match between the BtuC FP and the CGMD-FP (or HP, Fig. S3a), we did two sets of controls. First, we chose constructs at or near peaks in the FP and CGMD-FP and mutated multiple hydrophobic residues (Leu, Ile, Val, Met) located 40-50 residues from the PTC to less hydrophobic Ala residues (Fig. S4). The mutations caused significant drops in *f*_*FL*_, (*p*<0.01, two-sided t-test), except for construct *N*=191 that is mutated at the extreme N-terminus of TMH5. The mutation data allowed us to identify the membrane integration of TMHs 1, 2, 3, 4, 5, 7, 8, 9, and 10 with peaks I, II, III, IV, V, VIII, IX, X, and XI, respectively; the overlapping peaks VIII and IX represent the concerted integration of the closely spaced TMH7 and TMH8. However, peak II is shifted to unexpectedly high, and peaks X and XI to unexpectedly low, *N*_*start*_ values, Fig. 4c. To confirm these assignments, we obtained FPs for the isolated TMH2 (dashed green), TMH8 (dashed blue), and TMH10 (dashed pink) sequences, Fig. 4b,d; the individual FPs overlap the corresponding peaks II, IX, and XI in the full FP.

**Figure 4.**
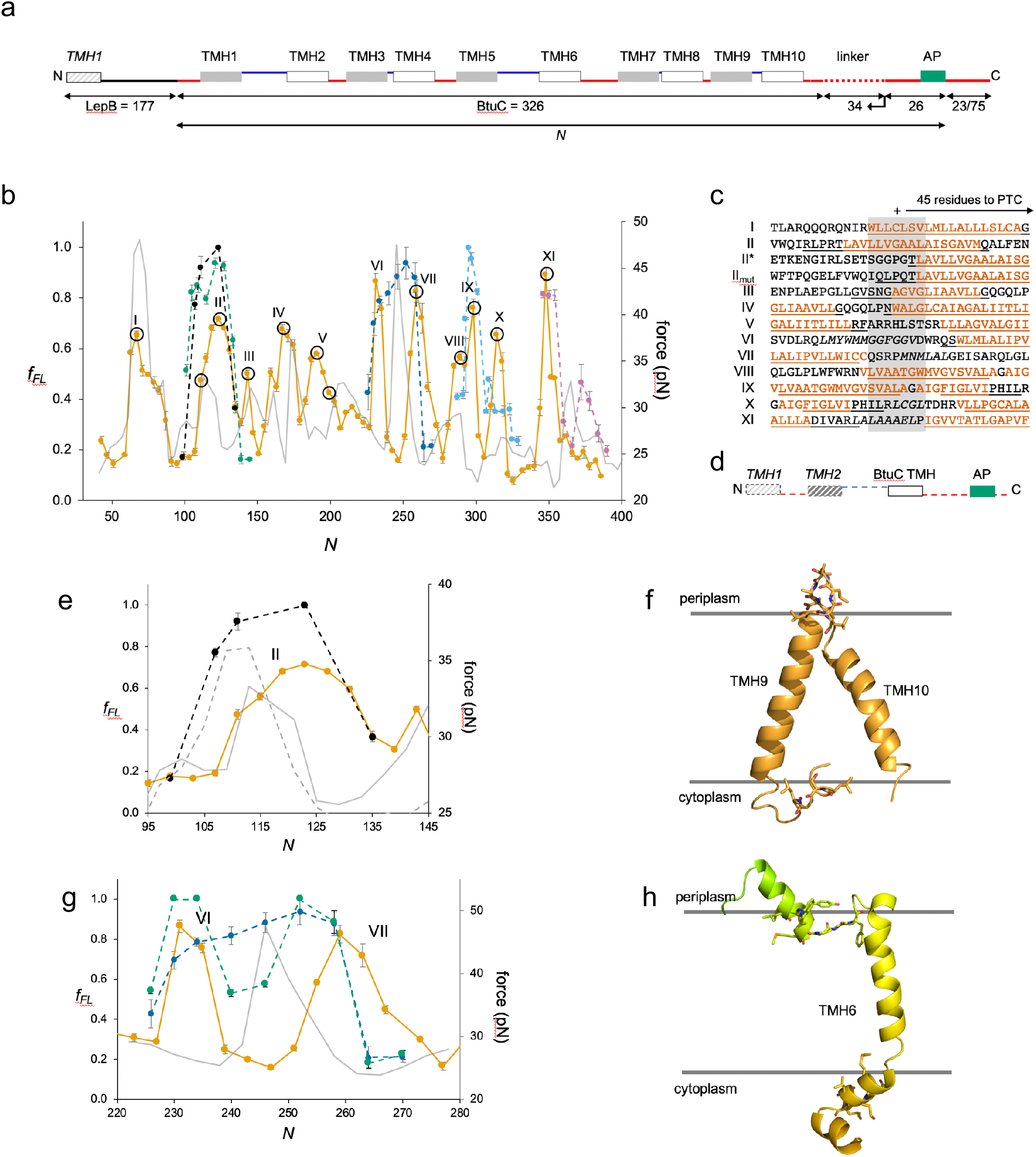
BtuC. (a) Construct design, c.f., Fig. 2a. The N-terminal LepB fusion is indicated. For constructs with *N*≥298, the C-terminal tail is 75 residues long. Circles indicate constructs for which mutations were made in the corresponding TMH, see Fig. S4. (b) FPs for BtuC (orange), BtuC-TMH2 (green), BtuC(R^47^R^56^R^59^→QQQ) (black), BtuC-TMH6 (dark blue), BtuC-TMH8 (blue), BtuC-TMH10 (pink), and CGMD-FP calculated with a −100 mV membrane potential (grey). Error bars indicate SEM values. (c) Sequences corresponding to peaks I-XI aligned from their *N*_*start*_ values. Hydrophobic TMH segments shown in orange and membrane-embedded α-helices according to the OPM database (*32*) underlined. Re-entrant loops and surface helices discussed in the text are italicized. (d) Construct design for obtaining FPs of isolated N-out oriented BtuC TMHs. Dashed segments are derived from LepB. (e) Enlarged FPs for BtuC (orange) and (R^47^R^56^R^59^→QQQ) (black), together with CGMD-FPs calculated with (grey) and without (dashed grey) a −100 mV potential. (f) BtuC TMH9-TMH10, with hydrophobic flanking residues in stick representation (PDB ID: 2QI9). (g) Enlarged FPs for BtuC (orange), isolated TMH6 (residues 187-206; blue), and isolated TMH5-6 (residues 138-206; green). In the latter construct, LepB TMH2 was not included in order to maintain the correct membrane topology of the BtuC TMH5-TMH6 part. The CGMD-FP is in grey. (h) Structure of TMH6 including the upstream periplasmic re-entrant helix and the downstream cytoplasmic surface helix, with hydrophobic flanking residues in stick representation.

Remarkably, the N-terminal end of the isolated TMH2 is ∼45 residues away from the PTC at *N*_*start*_, suggesting that upstream sequence elements present in full-length BtuC delay the integration of TMH2 by ∼10 residues (compare II* and II in Fig. 4c). The most conspicuous feature in the upstream region of TMH2 is the presence of 3 positively charged Arg residues, an uncommon occurrence in a periplasmic loop (*24*). Indeed, when these are replaced by uncharged Gln residues in full-length BtuC, peak II (dashed black in Fig. 4b,e) becomes almost identical to the FP for the isolated TMH2; a similar behavior is seen when the CGMD-FP simulation is run without an electrical membrane potential, Fig. 4e. Upstream positively charged residues thus delay the membrane integration of the N_out_-oriented TMH2, possibly because of the energetic cost of translocating them against the membrane potential (*3*), or because they are temporarily retained in the negatively charged exit tunnel (*15*). The low *N*_*start*_ values for peaks X and XI are likely caused by the short upstream hydrophobic segments LCGL and LAAALEL (Fig. 4c,f), similar to peak III in GlpG.

Neither peak VI nor VII seem to represent the integration of TMH6, but instead flank the location expected from the CGMD-FP and HP and apparently correspond, respectively, to the membrane insertion of a short periplasmic re-entrant helix and of a short cytoplasmic surface helix, Fig. 4c,h. Mutation of three hydrophobic residues to Ala in the latter significantly reduces the amplitude of peak VII (Fig. S4, construct *N*=259). Further, the FP for the isolated TMH6, Fig. 4b,g (dashed dark blue) peaks in the location expected from the CGMD-FP, between peaks VI and VII, and the FP for the isolated TMH5-6 part that includes the re-entrant helix but lacks the downstream surface helix is intermediate between the full-length BtuC and the TMH6 FPs, Fig. 4g (dashed green). Thus, the membrane interactions of the periplasmic re-entrant helix and the cytoplasmic surface helix exert a strong effect on the membrane integration of the intervening TMH6.

The high-resolution view of the cotranslational integration of multi-spanning membrane proteins provided here show that translocating nascent chains experience a distinct transition to a more hydrophobic environment at a distance of ∼45 residues from the PTC, generating an oscillating force on the nascent chain that is ultimately transmitted to the PTC and varies in step with the appearance of each TMH in the vicinity of the SecYEG translocon channel. It seems likely that such oscillations can have multiple effects on the translation of membrane proteins, as recently demonstrated for ribosomal frameshifting (*25*), and may affect protein quality control (*26*). Charged residues and membrane-interacting segments such as re-entrant loops and surface helices flanking a TMH can advance, delay, or attenuate its interaction with the translocon and surrounding lipid. Point mutations in an upstream TMH can affect the pulling force generated by downstream TMHs in a highly position-dependent manner, suggestive of residue-specific interactions between TMHs during the membrane-integration process. Complementing *in vitro* unfolding/folding studies (*27, 28*), real-time FRET analyses (*15*), chemical crosslinking (*29*), structure determination (*30*), and computational modeling (*31*), high-resolution *in vivo* FPA can help identify the molecular interactions underlying cotranslational membrane protein biogenesis with single-residue precision.

## Acknowledgements

This work was supported by grants from the Knut and Alice Wallenberg Foundation (2012.0282), the Novo Nordisk Fund (NNF18OC0032828), and the Swedish Research Council (621-2014-3713) to GvH, and from NIGMS, National Institutes of Health, (R01GM125063) to TFM and MZ. This work used the Extreme Science and Engineering Discovery Environment (XSEDE) Bridges computer at PSC through allocation TG-MCB160013. XSEDE is supported by National Science Foundation grant number ACI-1548562. We thank Dr. Rickard Hedman (Stockholm University) for programming and maintenance of the EasyQuant software.

## Supplemental material

### Materials and methods

#### Enzymes and chemicals

All enzymes used in this study were purchased from Thermo Scientific and New England Biolabs. Oligonucleotides were from Eurofins Genomics. DNA isolation/purification kits and precast polyacrylamide gels were from Thermo Scientific. L-[^35^S]-methionine was obtained from PerkinElmer. Mouse monoclonal antibody against the HA antigen was purchased from BioLegend. Protein-G agarose beads were manufactured by Roche. All other reagents were from Sigma-Aldrich.

#### Cloning and Mutagenesis

##### EmrE

The previously described N_out_-C_out_ oriented EmrE(C_out_) version carrying mutations T^28^R, L^85^R and R^106^A was engineered in a pETDuet-1 vector (*13*). A series of constructs was designed by inserting nucleotides downstream of EmrE(C_out_) coding for a variable LepB-derived linker sequence (between 4 and 34 residues), the 9-residue long HA tag, the 17-residue long *E. coli* SecM AP, and a 23-residue long C-terminal tail. The following APs with stalling strenghts were used: SecM(*Ec*) (FSTPVWISQAQGIRAGP), SecM(*Ec*-*Ms*) (FSTPVWISQ<u>HAP</u>IR<u>GS</u>P, mutations underlined), and SecM(*Ec*-Sup1) (FSTPVWISQA<u>PP</u>IRAGP, mutations underlined). The LepB-derived linker as well as EmrE(C_out_) were truncated 2-5 residues at a time from the C-terminus of the respective sequence. Site-specific DNA mutagenesis was carried out to introduce point mutations E^14^L, E^14^A, E^14^D, and E^14^Q in EmrE(C_out_). All cloning and mutagenesis products were confirmed by DNA sequencing. Different EmrE sequences used in this study are summarized in Supplemental Table 1.

##### GlpG

The gene encoding for GlpG was amplified from the genome of *E. coli* K-12 MG1655 strain by PCR and assembled together with other sequence elements into pING1 plasmid by Gibson assembly®. For the longest truncates, a LepB-derived unstructured linker was introduced downstream of the GlpG sequence, followed by an HA tag, a 17-residue long *E. coli* SecM AP and a 23-residue long C-terminal tail derived from LepB. Partially overlapping primers were used in around-the horn PCR (https://www.protocols.io/view/around-the-horn-pcr-and-cloning-rf2d3qe) to create deletion variants truncating upstream of the HA-tag. All the sequences of GlpG deletion variants used in this study are summarized in Supplemental Table 1. For Lep-GlpG constructs, 60 N-terminal residues corresponding to the soluble domain were truncated from GlpG and substituted by LepB N-terminal segment comprising TMH1 and a long cytoplasmic loop (1-174 res of LepB). Three different stalling sequences of increasing strength were used: SecM(*Ec*) (FSTPVWISQAQGIRAGP), SecM(*Ec*-*Ms*) (FSTPVWISQ<u>HAP</u>IR<u>GS</u>P, mutations underlined), and SecM(*Ec*-Sup1) (FSTPVWISQA<u>PP</u>IRAGP, mutations underlined). Mutations in SecM(*Ec*) AP and GlpG folding variants NTD(F^16^E), GlpG(Y^138^F^139^L^143^→NNN) were engineered using partially overlapping primers in around-the-horn PCR. All cloning and mutagenesis products were confirmed by DNA sequencing.

For *in vitro* transcription/translation of the soluble NTD domain, constructs of variable length were fused to the SecM(*Ec*) AP and cloned into the pET19b vector. Folding variant NTD(F^16^E) was engineered using partially overlapping primers in around-the-horn PCR. pET19b plasmids containing different GlpG variants were used as template to create linear DNA fragments amplified by PCR for each construct using forward and reverse primers that anneal to the T7 promoter and terminator regions, respectively.

##### BtuC

The previously described pING1 plasmid harboring a truncated LepB sequence with an inserted hydrophobic test segment (6L/13A) followed by a variable LepB-derived linker (between 9 and 43 residues), the 17-residue long *E. coli* SecM AP, and a C-terminal tail comprised of 23 or 75 residues derived from LepB was used to generate all BtuC constructs (*4*). All BtuC sequences used in this study are summarized in Table S1.The gene encoding BtuC was amplified from the genome of the *E. coli* K-12 MG1655 strain by PCR and then engineered to replace 6L/13A using Gibson assembly^**®**^ (*33*). In order to maintain the correct topology of BtuC, the sequence coding for TMH2 of LepB (between residues P^58^ and P^114^) was removed by deletion-PCR resulting in a 177-residue long sequence upstream of BtuC. A gene sequence encoding 52 residues (part of LepB P2 domain) was introduced downstream of the SecM AP for constructs with *N*≥298, resulting in an extension of the C-terminal tail from 23 to 75 residues in order to improve protein separation during SDS-PAGE. The LepB-derived linker as well as BtuC were truncated 4 residues at a time from the C terminus of the respective sequence. Site-specific DNA mutagenesis was carried out to replace 3 or 6 hydrophobic residues with Ala residues in TMHs of BtuC and to replace 3 Arg residues with Gln residues in the periplasmic loop connecting TMH1 and TMH2 (R^47^R^56^R^59^→QQQ). Gene sequences of single TMH and 2-TMH constructs were cloned with the variable linker sequence derived from LepB, the single TMH constructs were placed in the background containing gene sequences of both LepB TMHs in order to maintain the correct topology. Furthermore, BtuCΔLepB constructs lacking the N-terminal LepB fusion were obtained by deletion of the entire LepB sequence upstream of BtuC, and the 9-residue long LepB-derived linker was replaced with an HA tag for immunoprecipitation. All cloning and mutagenesis products were confirmed by DNA sequencing.

#### In vivo pulse-labeling analysis

Competent *E. coli* MC1061 or BL21 (DE3) cells were transformed with the respective pING1 (BtuC, GlpG) or pET Duet-1 (EmrE) plasmid, respectively and grown overnight at 37°C in M9 minimal medium supplemented with 19 amino acids (1 µg/ml, no Met), 100 μg/ml thiamine, 0.4% (w/v) fructose, 100 mg/ml ampicillin, 2 mM MgSO_4_, and 0.1mM CaCl_2_. Cells were diluted into fresh M9 medium to an OD_600_ of 0.1 and grown until an OD_600_ of 0.3 −0.5. Expression from pING1 was induced with 0.2% (w/v) arabinose and continued for 5 min at 37°C. Expression from pET Duet-1 was induced with 1 mM IPTG and continued for 10 min at 37°C. Proteins were then radiolabeled with [^35^S]-methionine for 2 min (1 min for BtuC constructs lacking the N-terminal LepB fusion) at 37°C before the reaction was stopped by adding ice-cold trichloroacetic acid (TCA) to a final concentration of 10%. Samples were put on ice for 30 min and precipitates were spun down for 10 min at 20,000 g at 4°C in a tabletop centrifuge (Eppendorf). After one wash with ice-cold acetone, centrifugation was repeated and pellets were subsequently solubilized in Tris-SDS buffer (10 mM Tris-Cl pH 7.5, 2% (w/v) SDS) for 5 min while shaking at 1,400 rpm at 37°C. Samples were centrifuged for 5 min at 20,000 g to remove insoluble material. The supernatant was then added to a buffer containing 50 mM Tris-HCl pH 8.0, 150 mM NaCl, 0.1 mM EDTA-KOH, 2% (v/v) triton X-100, and supplemented with Pansorbin^®^ (BtuC constructs) or Gammabind G Sepharose^®^ (all GlpG and EmrE constructs, and BtuC constructs lacking the N-terminal LepB fusion). After 15 min incubation on ice, non-specifically bound proteins were removed by centrifugation at 20,000xg (when Pansorbin^®^ was used) or 7,000xg (when Gammabind G Sepharose^®^ was used). The supernatant was used for immunoprecipitation of BtuC constructs using Pansorbin® and LepB antisera (rabbit), or immunoprecipitation of GlpG/EmrE constructs using Gammabind G Sepharose^®^ and Anti-HA.11 Epitope Tag Antibody (mouse). The incubation was carried out at 4°C whilst rolling. After centrifugation for 1 min, immunoprecipitates were washed with 10 mM Tris-Cl pH 7.5, 150 mM NaCl, 2 mM EDTA, and 0.2% (v/v) triton X-100 and subsequently with 10 mM Tris-Cl pH 7.5. Samples were spun down again and pellets were solubilized in SDS sample buffer (67 mM Tris, 33% (w/v) SDS, 0.012% (w/v) bromophenol blue, 10 mM EDTA-KOH pH 8.0, 6.75% (v/v) glycerol, 100 mM DTT) for 10 min while shaking at 1,400 rpm. Solubilized proteins were incubated with 0.25 mg/ml RNase for 30 min at 37°C and subsequently separated by SDS-PAGE. Gels were fixed in 30% (v/v) methanol and 10% (v/v) acetic acid and dried by using a Bio-Rad gel dryer model 583. Radiolabeled proteins were detected by exposing dried gels to phosphorimaging plates, which were scanned in a Fuji FLA-3000 scanner. Band intensity profiles were obtained using the FIJI (ImageJ) software and quantified with our in-house software EasyQuant. Data was collected from three independent biological replicates, and averages and standard errors of the mean (SEM) were calculated.

#### In vitro transcription/translation of GlpG NTD

*In vitro* transcription/translation was performed using the commercially available PURExpress system (New England Biolabs). Reactions were mixed according to the manufacturer’s recommendations by the addition of 2.2 μl of linear DNA of each construct giving a final volume of 10 μl. Polypeptide synthesis was carried out in the presence of [^35^S]-methionine at 37°C for 15 min under 700 rpm shaking. Translation was stopped by the addition of trichloroacetic acid (TCA) to a final concentration of 5% and incubated on ice for at least 30 min. Total protein was sedimented by centrifugation at 20,000 g for 10 min at 4°C in a tabletop centrifuge (Eppendorf). The pellet was resuspended in 2x SDS/PAGE sample buffer, supplemented with RNaseA (400 µg/ml) to digest the stalled peptidyl-tRNA, and incubated at 37°C for 15 min under 1,000 rpm agitation. The samples were resolved on 12% Bis-Tris gels (Thermo Scientific) in MOPS buffer. Gels were dried (Hoefer GD 2000), exposed to a phosphorimager screen for 24 h and scanned using the Fujifilm FLA-9000 phosphorimager for visualization of radioactively labeled protein species.

#### Molecular dynamics simulations

Computer simulations of cotranslational membrane integration were carried out using a previously developed and validated coarse-grained molecular dynamics (CGMD) model in which nascent proteins are mapped on to CG beads representing three amino acids (*10, 34*). The nascent protein interacts with the Sec translocon and the ribosome via pairwise interactions that depend on the hydrophobicity and charge of the beads of the nascent protein. The interaction parameters are unchanged from previous work (*34*). The lateral gate of the translocon switches between the open and closed conformations with probability dependent on the difference in free energy between the two conformations. The structures of the ribosome and translocon are based on cryo-EM structures and, aside from the lateral gate of the translocon, are fixed in place during the simulations. The lipid bilayer and cytosol are modelled implicitly. The positions of the nascent protein beads are evolved using overdamped Langevin dynamics with a timestep of 300 ns and a diffusion coefficient of 253 nm^2^/s. Membrane potentials are included by adding an electrostatic energy term to the simulations, as previously described (*10*).

To simulate protein translation, new amino acids are added to the nascent chain at a rate of 5 amino acids per second. Simulations of EmrE, GlpG, and BtuC begin with 12 amino acids translated. Translation continues until the nascent protein reaches the desired length, at which point translation is halted and forces on the C-terminus of the nascent chain are measured every 3 ms for 6 seconds. This methodology has been found to accurately reproduce experimental force profiles (*10*). Forces are measured starting at a nascent protein length of 18 amino acids for EmrE and BtuC, and 70 for GlpG. The computational force profile (CGMD-FP) is then obtained by measuring the forces at lengths incremented by four amino acids. Simulations at different lengths are performed independently and repeated 100 times. Because the ribosomal exit tunnel is truncated in the CGMD model, a shift in the protein index is required to compare simulated and experimental results. Shifts of −12, −5, and −5 residues are used for EmrE, GlpG and BtuC CGMD-FPs, respectively. The shifts are estimated by aligning the computational and experimental force profiles and are in line with what is expected given the length of the truncated exit tunnel. Variation in the shift may reflect different degrees of compaction of the nascent chain. Although previous work provides a framework to estimate the experimentally observed fraction full-length from simulated forces given a specific arrest peptide (*10, 35*), forces are reported directly to facilitate comparison between experiments performed with different arrest peptides.

## Abbreviations

FRET: fluorescence resonance energy transfer
AP: arrest peptide
PTC: peptidyl transferase center
FPA: force profile analysis
CGMD-FP: coarse-grained molecular dynamics force profile
HP: hydrophobicity plot
NTD: N-terminal domain
TMH: transmembrane helix

---

**Fig. S1.**
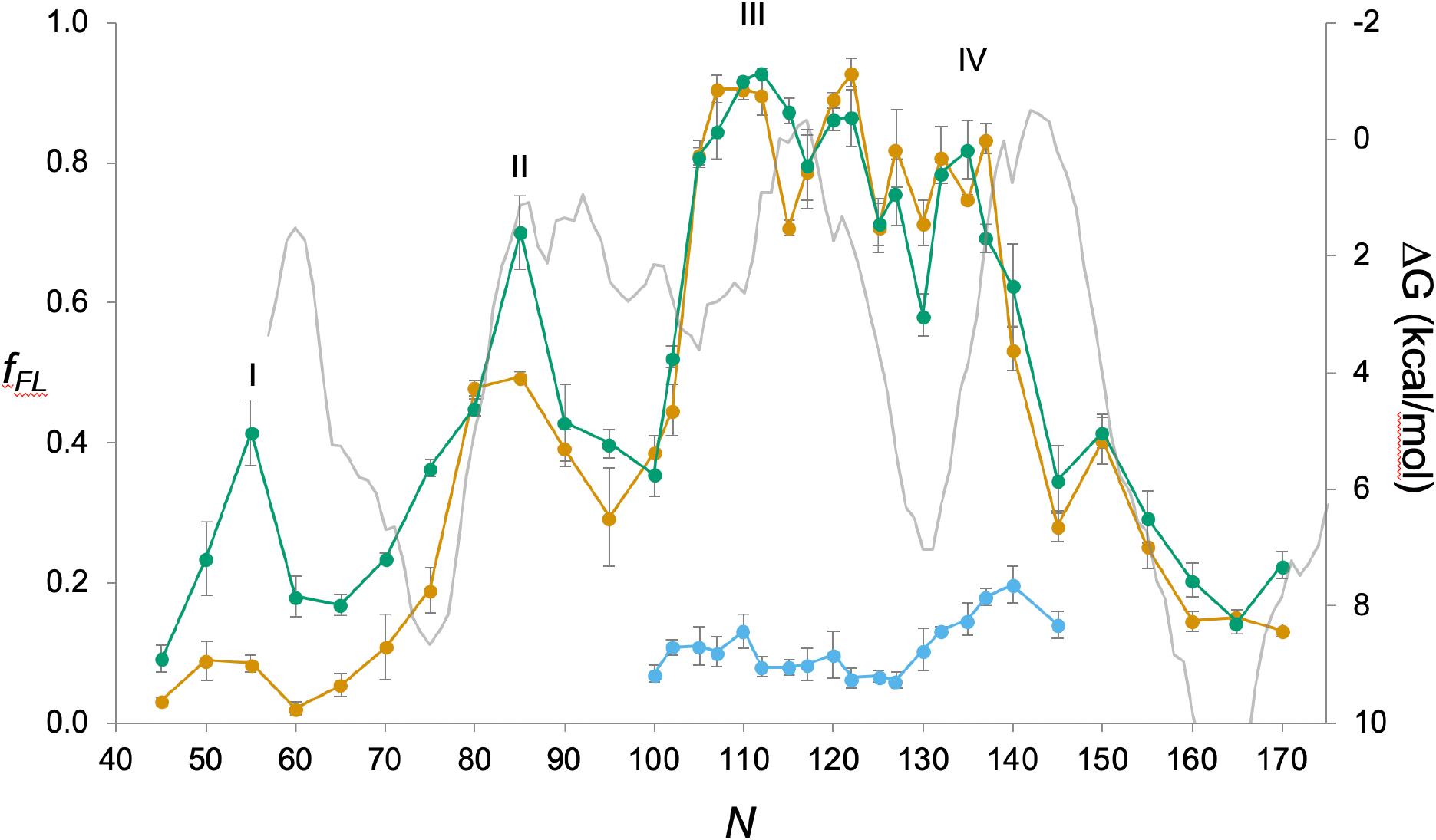
EmrE(C_out_). As in Fig. 2b, but with a HP (ΔG) calculated by TOPCONS (*2, 36*) (grey). Since the HP represents the membrane-integration energy and the FP the force generated during integration, the two profiles have been aligned such that peaks in the FP approximately align with maxima in the derivative of the HP.

**Fig. S2.**
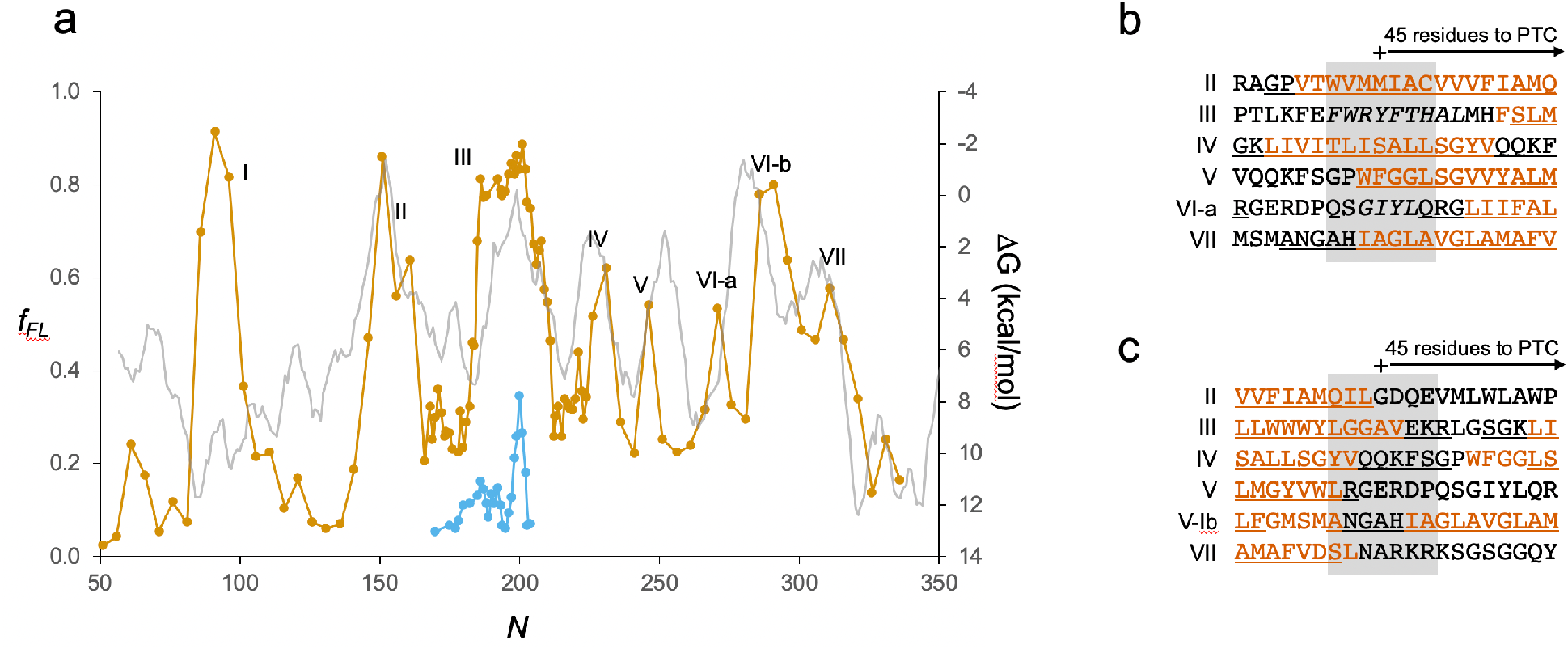
GlpG. (a) As in Fig. 3b, but with a HP (ΔG) calculated by TOPCONS (*2, 36*) (grey). (b) Sequences corresponding to peaks II-VII aligned based on the *N*_*start*_ values. The periplasmic surface helix upstream of TMH2 and the hydrophobic patch upstream of TMH5 are in italics. Hydrophobic TMH segments shown in orange and membrane-embedded α-helices underlined. (c) Sequences corresponding to peaks II-VII aligned based on the *N*_*end*_ values. Hydrophobic TMH segments shown in orange and membrane-embedded α-helices underlined.

**Fig. S3.**
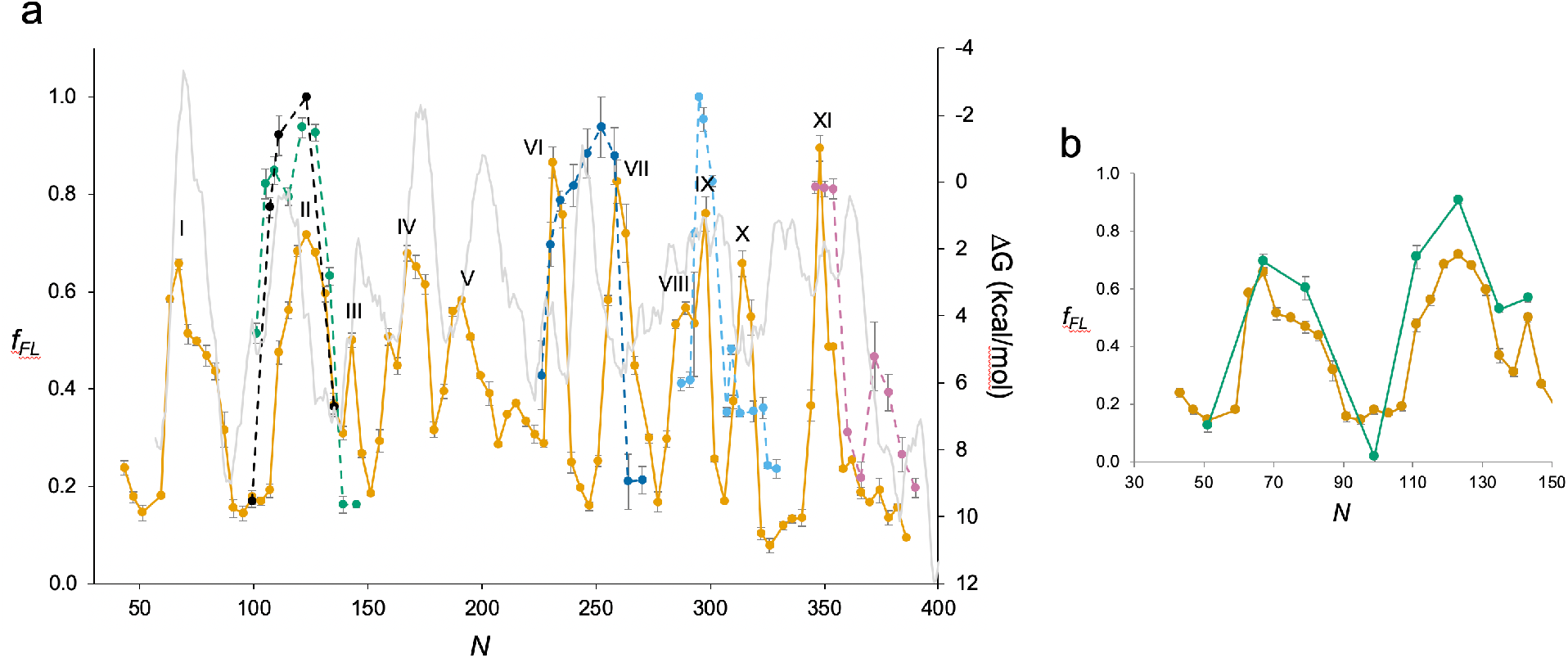
BtuC. (a) As in Fig. 4b, but with a HP (ΔG) calculated by TOPCONS (*2, 36*) (grey). (b) Close-up view of the BtuC FP (*N*=30-150; orange), and the corresponding FP obtained with GlpG constructs lacking the N-terminal LepB fusion (green).

**Fig. S4.**
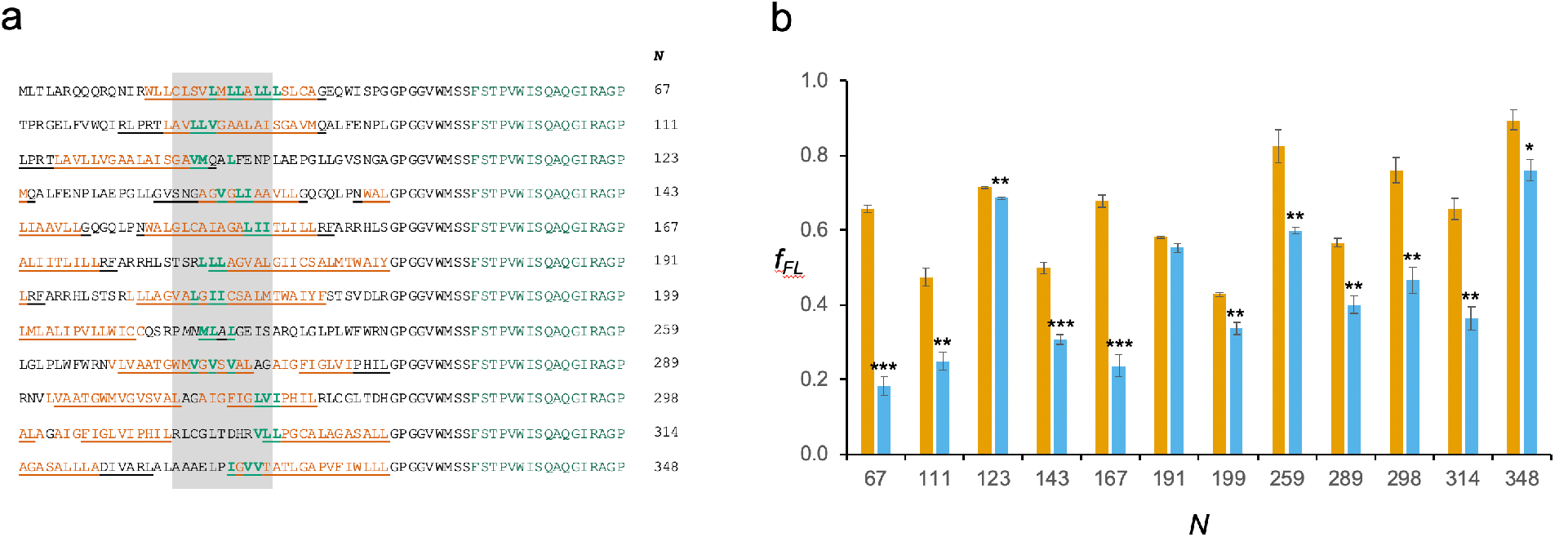
Mutations in constructs representing peaks in the BtuC FP. (a) Sequences of the 67 residues leading up to the end of the AP for constructs with the indicated *N*-values. The constructs are identified by black circles on the BtuC FP in Fig. 4b. For each construct, the residues indicated in bold green were simultaneously mutated to Ala. The shaded area encompasses residues located 40-50 residues away from the C-terminal end of the AP in the respective constructs. Hydrophobic TMH segments are shown in orange and membrane-embedded α-helices are underlined. (b) *f*_*FL*_ values for the unmutated constructs (orange) and the Ala-replacement mutants (blue). Error bars indicate SEM values, and stars indicate p-values calculated using a two-sided t-test (p < 0.05, *; p < 0.01, **; p < 0.001, ***).

**Supplementary Table S1.**
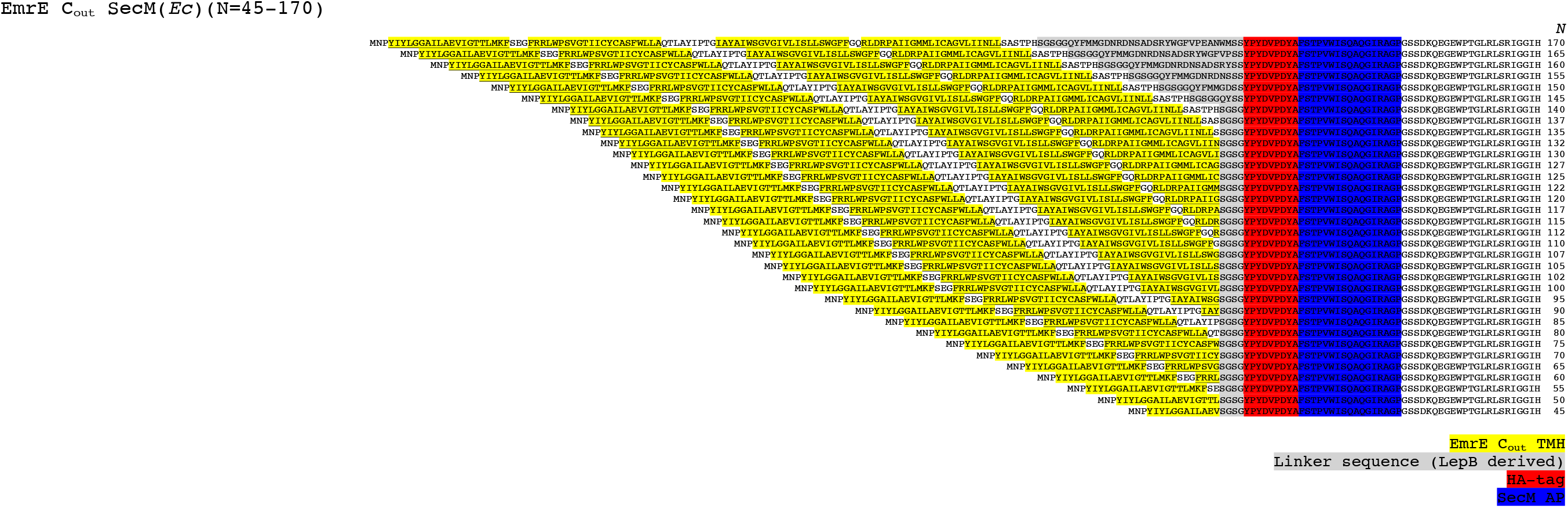
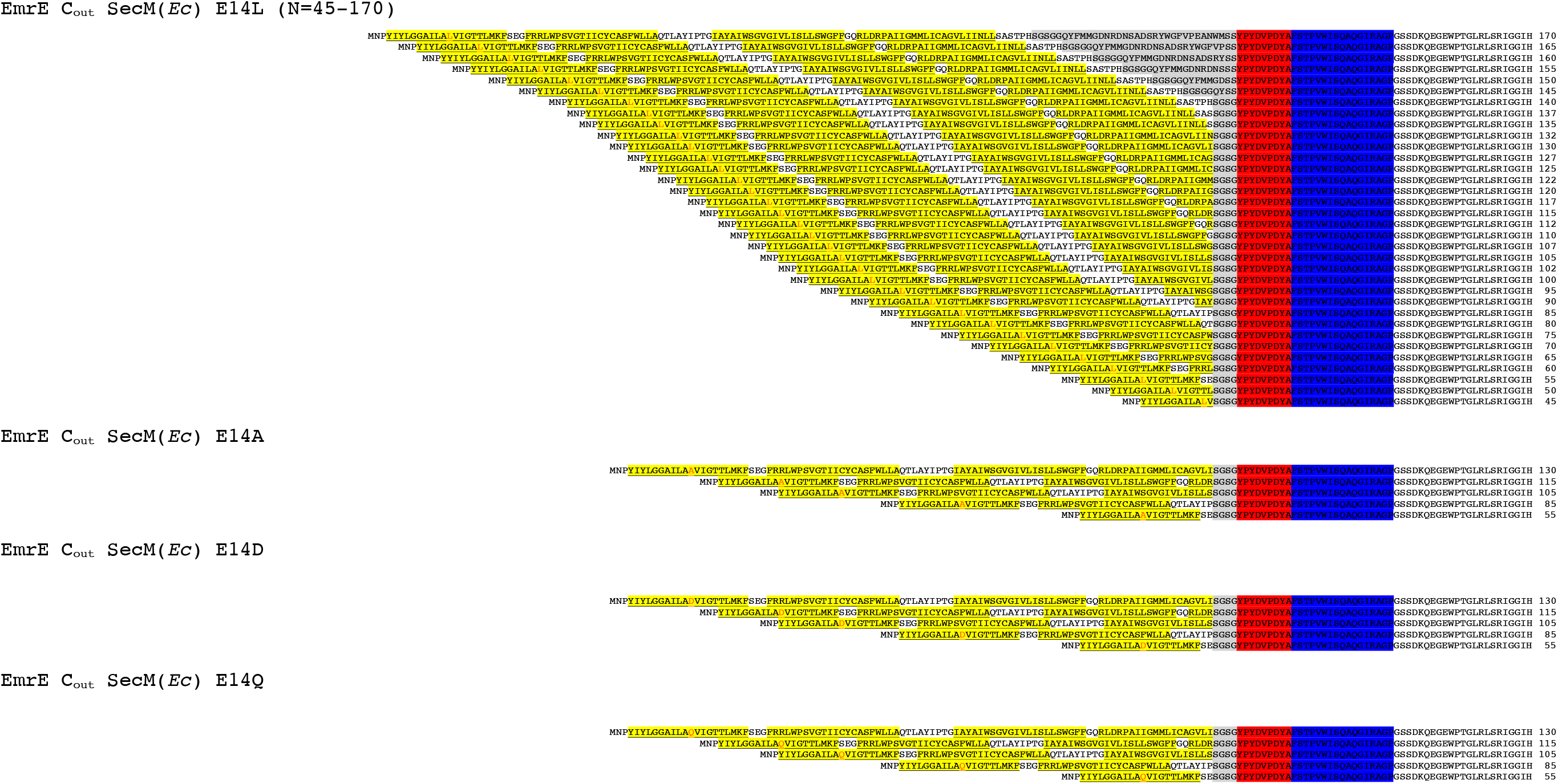
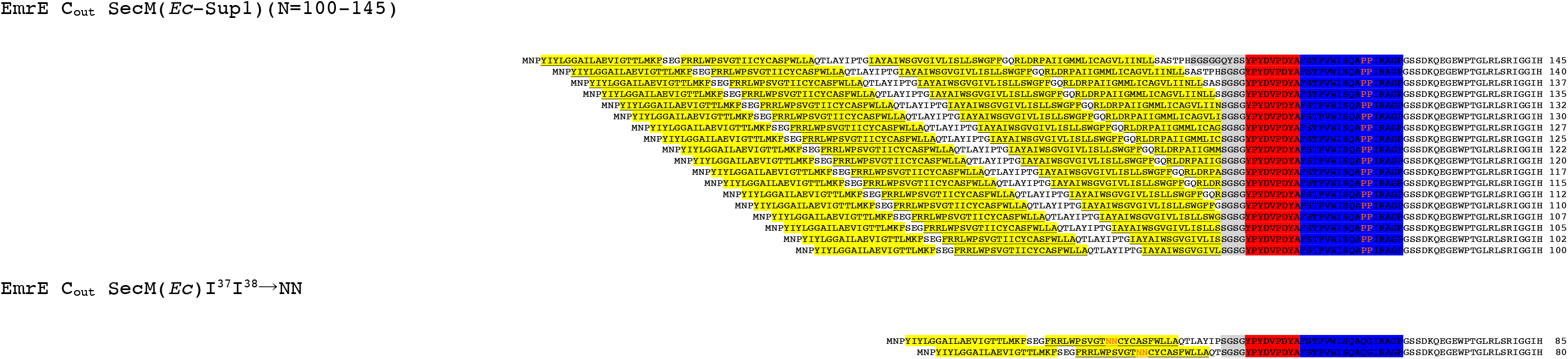
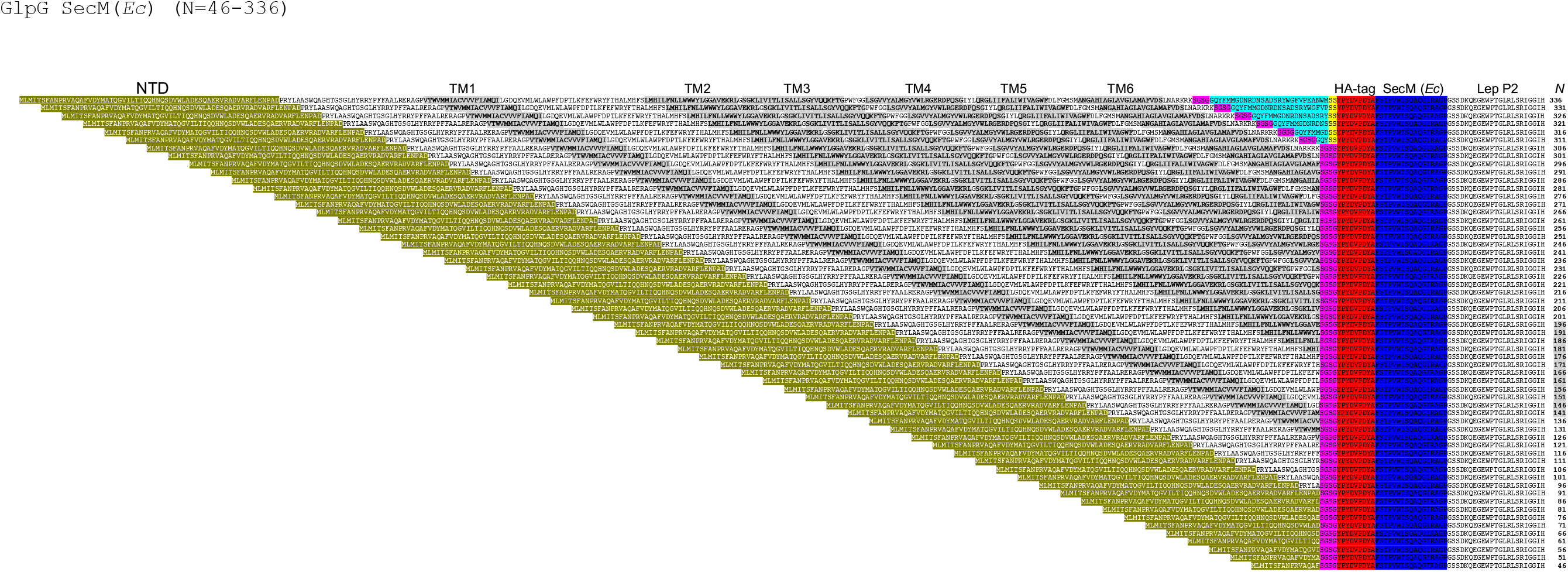
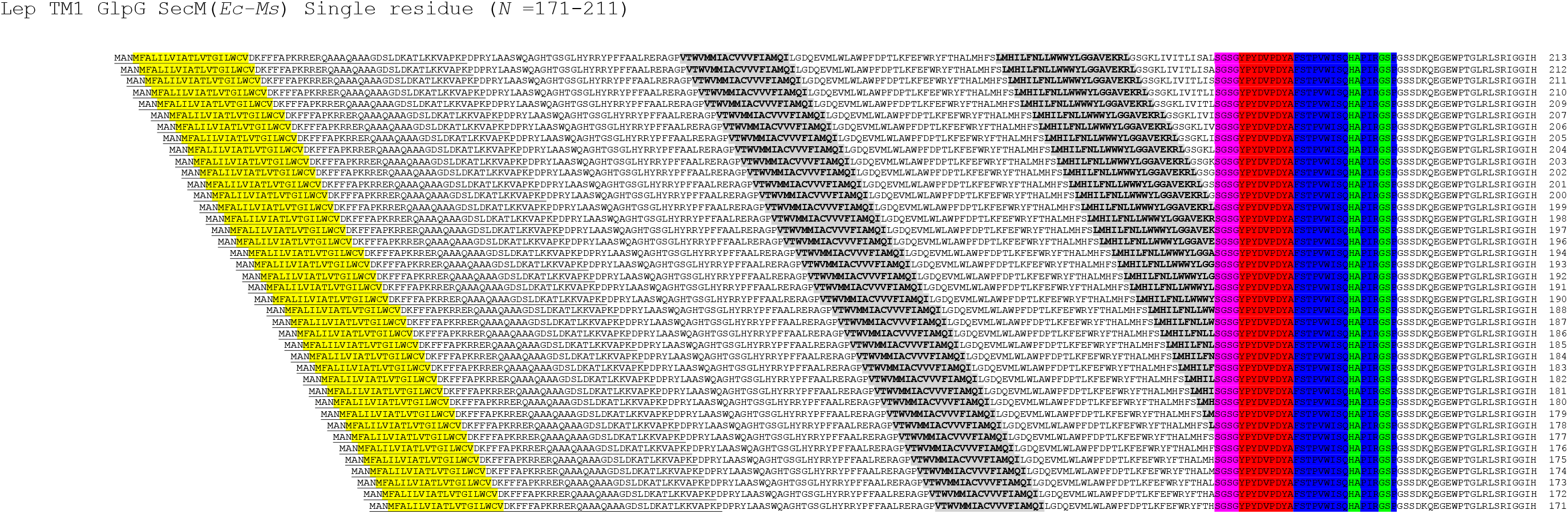
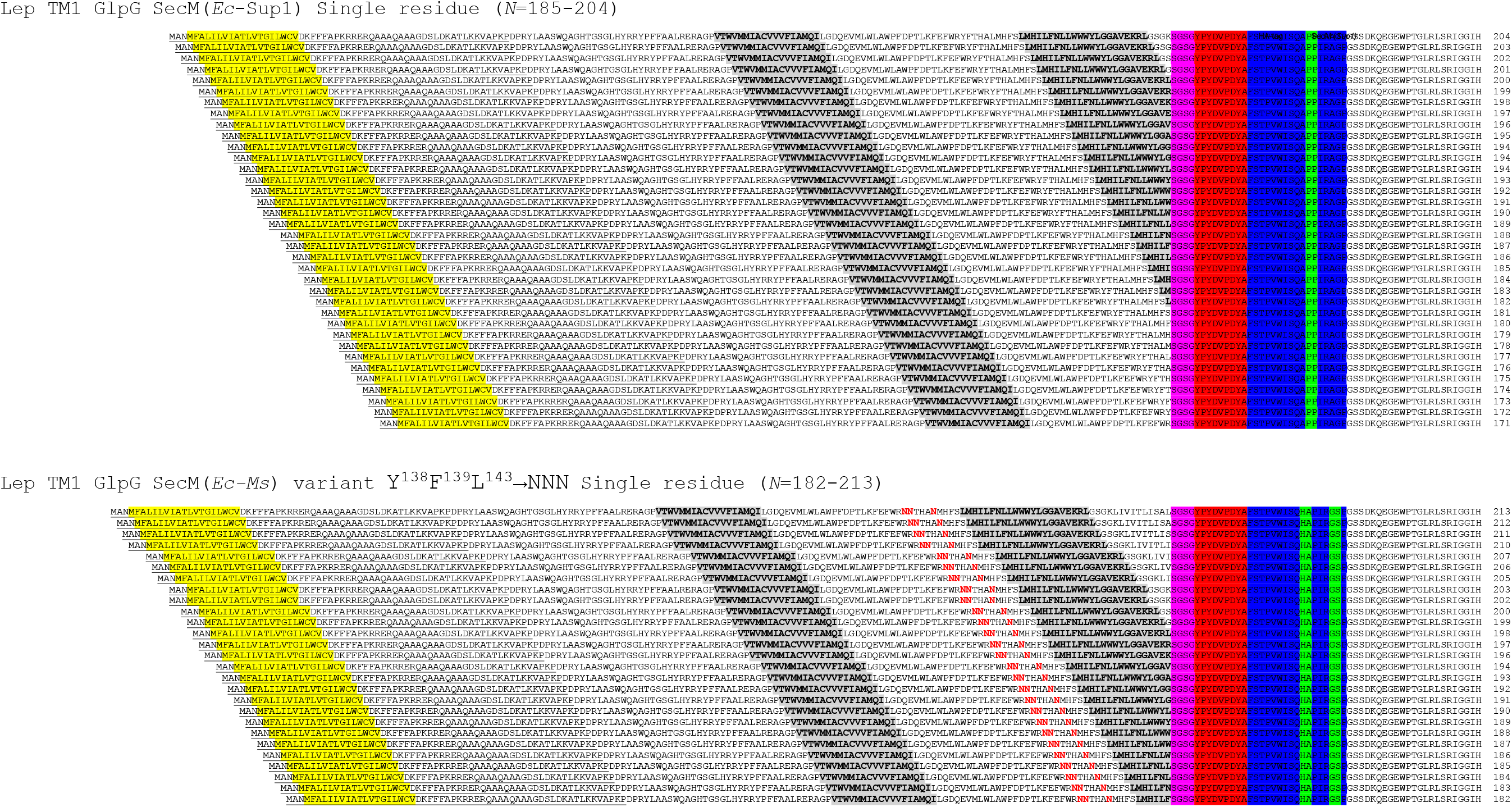
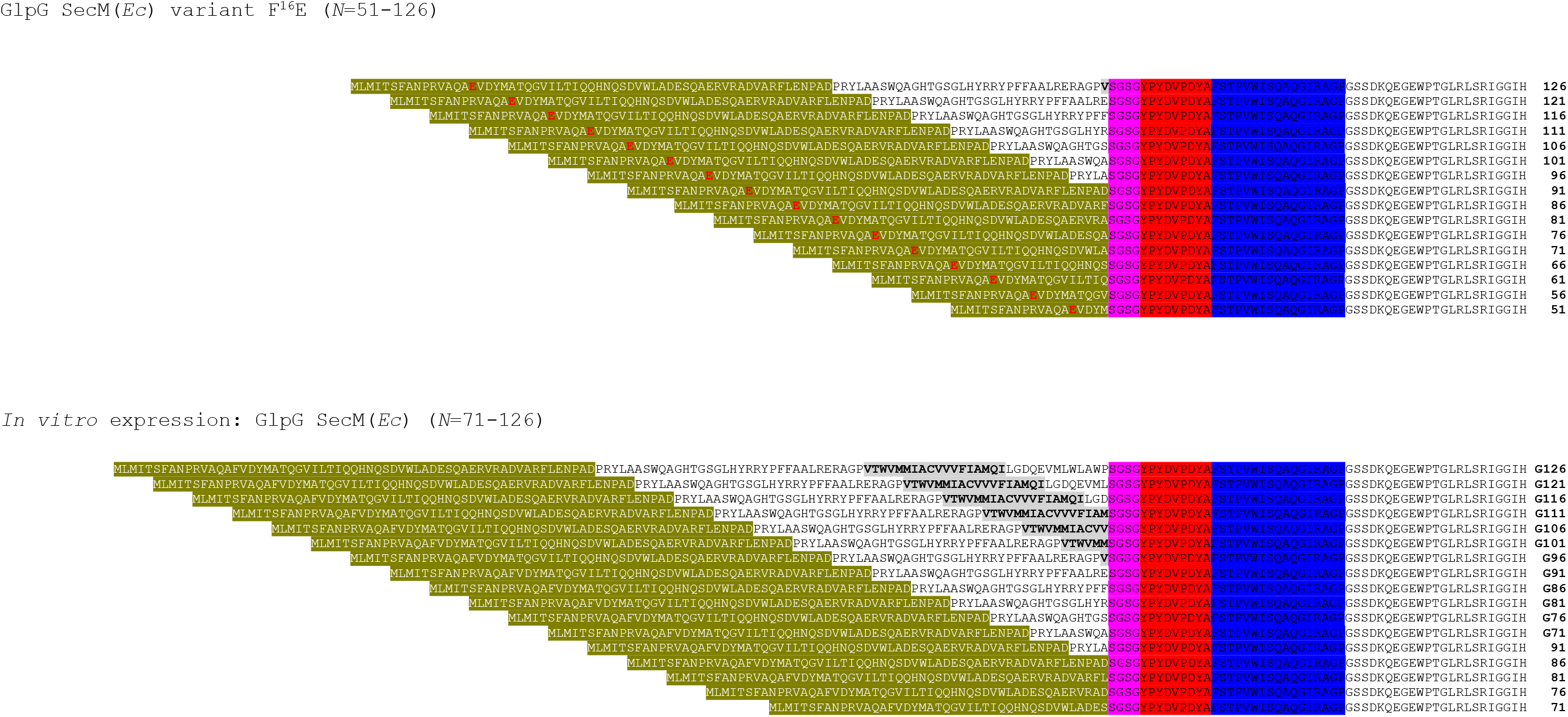
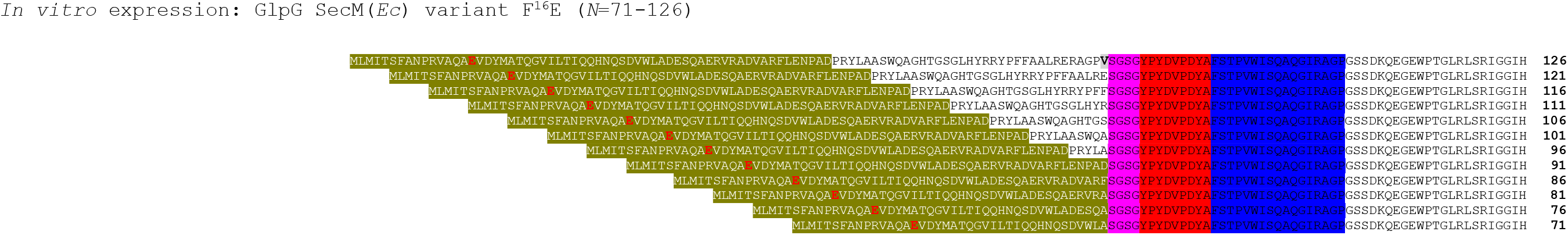
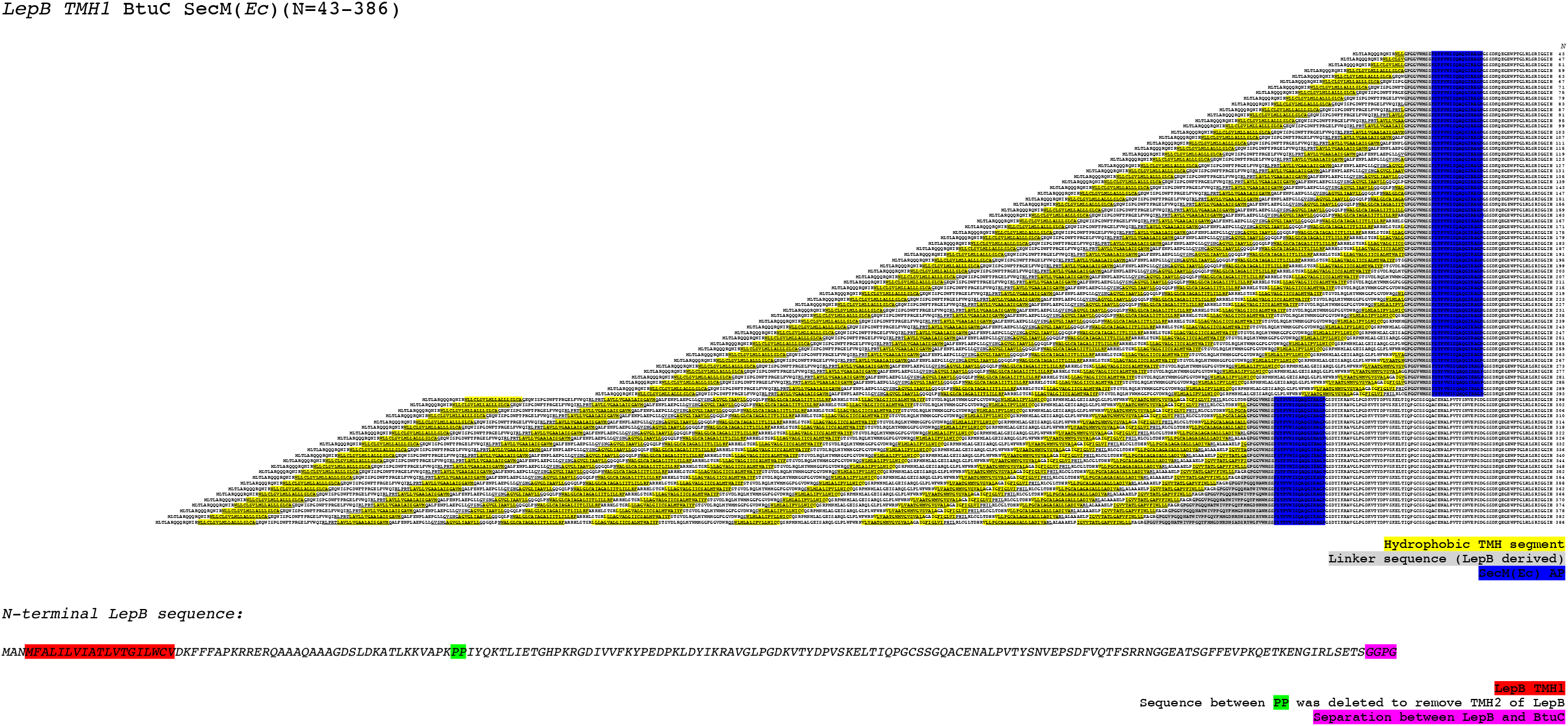
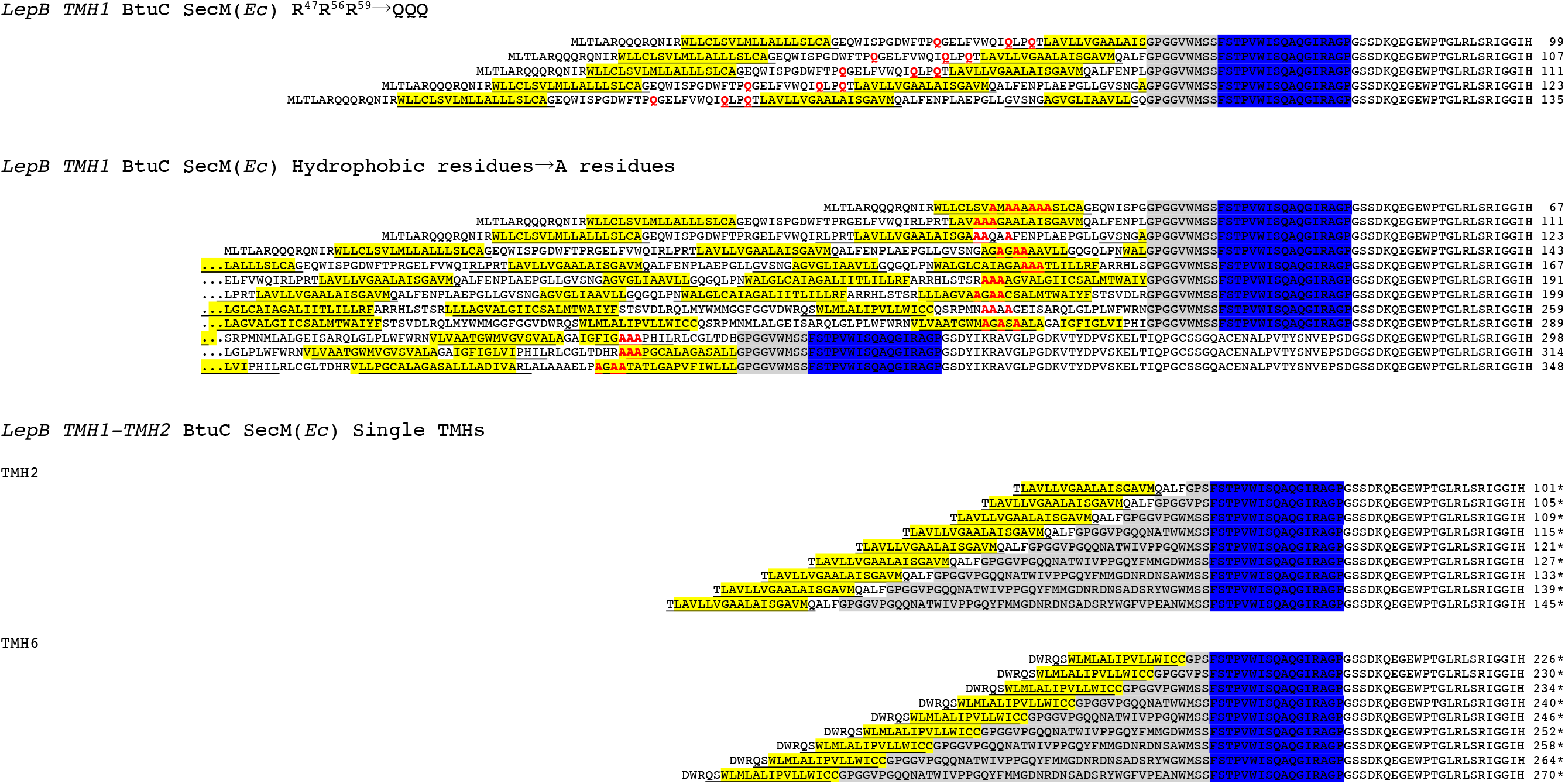
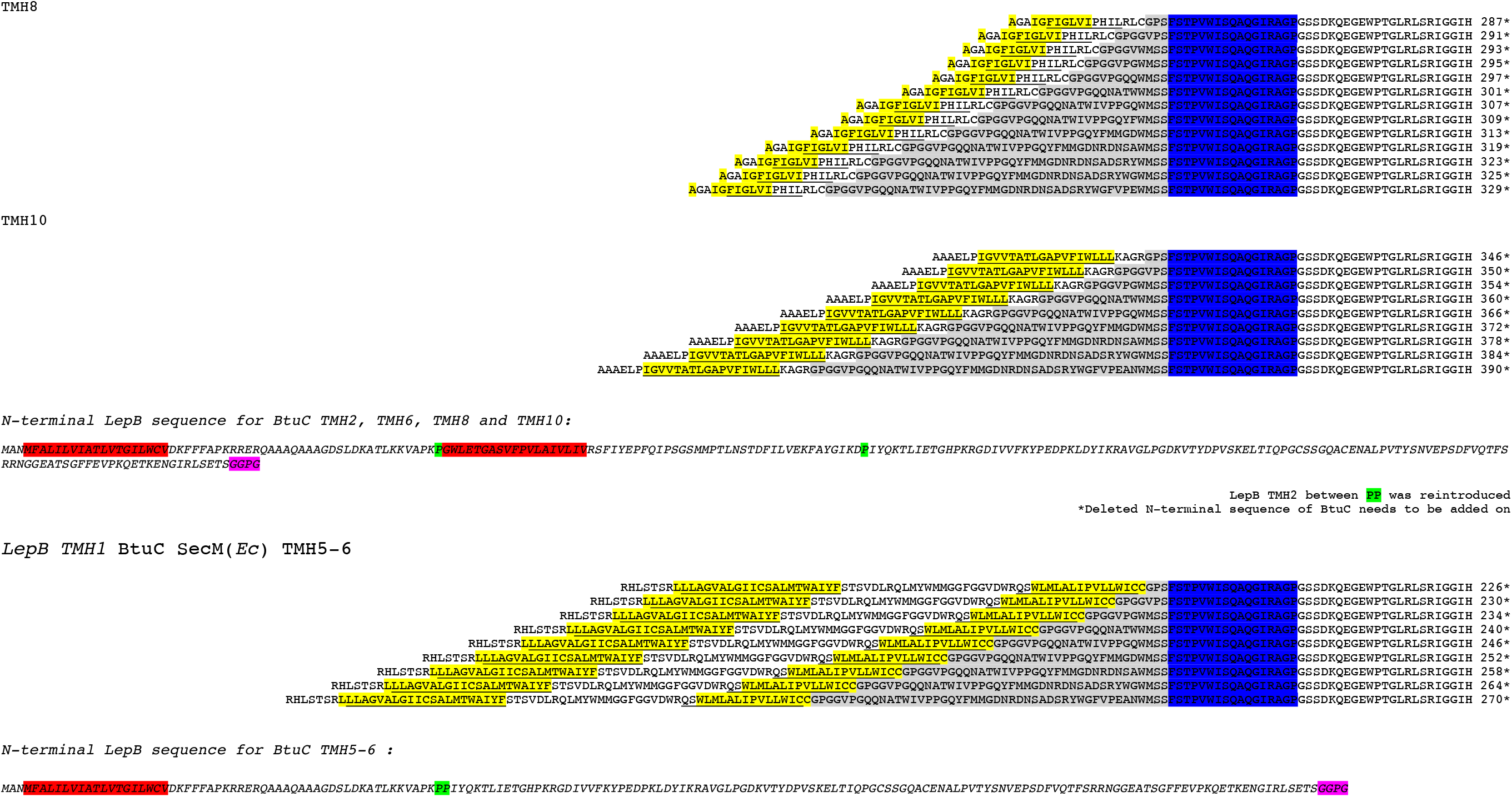
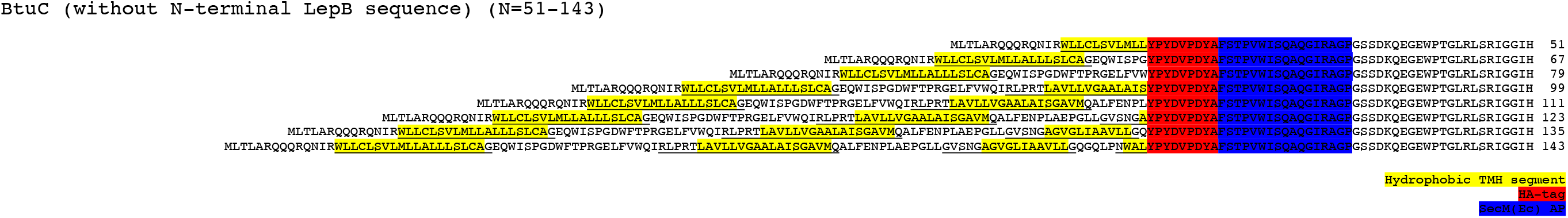
Amino acid sequences of EmrE(C_out_), GlpG, and BtuC constructs.

